# Altered Motor Awareness in Parkinson’s Disease with Progressive Micrographia

**DOI:** 10.64898/2026.01.29.702455

**Authors:** Hiroyuki Hamada, Kyohei Mikami, Wen Wen, Yoshihiro Itaguchi, Atsushi Yamashita, Qi An

## Abstract

Micrographia, characterized by abnormally small handwriting, is a common motor symptom in Parkinson’s disease (PD), and has been classified into consistent micrographia (CM) and progressive micrographia (PM), which may reflect different underlying mechanisms. However, the relative contributions of motor execution and sensorimotor processing to these subtypes remain unclear. In this study, forty-five people with Parkinson’s disease (PwP) and twenty age-matched healthy controls (HCs) completed a handwriting task and a control detection task (CDT) assessing the ability to distinguish self-generated from externally generated movements. PwP were classified based on handwriting metrics as showing no micrographia, CM, and/or PM. CDT accuracy was significantly lower in PwP than in HCs (*p* < 0.01) and was particularly reduced in PwP with PM (*p* < 0.01). These findings indicate that PM is associated with impaired motor awareness, suggesting altered sensorimotor integration as a characteristic feature of this subtype.

## Introduction

Parkinson’s disease (PD) is a progressive neurodegenerative disorder that primarily affects motor function. It is clinically characterized by core motor symptoms, including tremors, rigidity, bradykinesia, and postural instability. In addition to these motor manifestations, PD is associated with a range of non-motor symptoms, such as autonomic dysfunction and cognitive decline, that arise from the degeneration of dopamine-producing neurons in the substantia nigra pars compacta.^1^ Among the motor symptoms of people with Parkinson’s disease (PwP), micrographia is one of the most characteristic.^2^ It occurs in about 50% of PwP,^3^ and in severe cases, the handwriting becomes so small and distorted that it is difficult for both the individual and others to decipher, thereby posing significant challenges in daily activities such as signing documents or taking notes. Micrographia is typically classified into two subtypes: consistent micrographia (CM), in which the handwriting is small from the outset, and progressive micrographia (PM), where the size of characters gradually diminishes as writing continues.^4^ In both cases, the reduced legibility of handwriting has prompted the exploration of pharmacological and rehabilitative interventions. While pharmacological treatment has shown effectiveness in alleviating CM,^5^ PM remains particularly resistant to conventional medication and rehabilitation strategies,^6^ highlighting the need for the development of more effective intervention approaches, particularly for PM.

The differential responsiveness to intervention between the two subtypes of micrographia is thought to reflect differences in their underlying mechanisms. CM has been shown to be associated with disease severity, as measured by the Movement Disorder Society–Unified Parkinson’s Disease Rating Scale (MDS-UPDRS) motor score, and with bradykinesia.^7^ CM symptoms have also been reported to improve following administration of levodopa or deep brain stimulation.^4,8^ In contrast, PM does not appear to be significantly related to motor function,^9^ and pharmacological interventions have not been found to exert direct therapeutic effects.^10^ As such, the pathophysiological mechanisms underlying PM remain poorly understood, and no effective pharmacological or rehabilitative interventions have yet been established for its treatment.

Neurophysiological studies have revealed that in PwP presenting with CM, neurodegeneration is primarily localized to the basal ganglia. In contrast, PwP with PM exhibit not only degeneration in the basal ganglia, but also functional abnormalities in broader motor-related networks, including the supplementary motor area, cingulate motor area, and cerebellar motor regions.^11,12^ Furthermore, evidence that the size of written characters differs between eyes open and eyes closed conditions during the off-medication state in PD,^13^ along with findings that somatosensory-based awareness is essential for handwriting,^14^ suggests that micrographia may result, at least in part, from an impaired ability to correct discrepancies between motor predictions and sensory feedback.

Although research on sensory feedback processing in PwP remains limited, several studies have reported altered awareness of movement. Previous research has indicated cognitive alterations in motor control, such as changes in the awareness of movement timing and direction.^15^ These findings suggest a disruption in the sense of agency, which arises from the comparison between predicted motor outcomes and actual sensory feedback. While PwP typically exhibit mild or no overt sensory deficits, they may show impairments in the central processing mechanisms that integrate and compare sensory and motor information. However, the link between altered motor awareness and micrographia remains poorly understood.

Therefore, the present study aimed to clarify differences in awareness of motor control between healthy individuals and PwP, as well as to investigate the relationship between the characteristics of micrographia and changes in movement self-perception in PwP. Specifically, we employed a control detection task (CDT),^16^ a behavioral paradigm designed to distinguish self-generated from externally generated movements, to examine how CDT performance relates to micrographia subtypes, with the goal of elucidating potential mechanisms underlying PM.

The CDT is based on the theoretical framework of sensorimotor integration and predictive coding, and is commonly used to assess the sense of agency by requiring participants to identify which of two stimuli they are controlling. In this task, participants operate a cursor or object under variable visual feedback and must discriminate whether the observed movement corresponds to their own motor output or to an externally generated trajectory. This paradigm quantifies the correspondence between predicted motor outcomes and actual sensory feedback, providing a behavioral index of action–outcome monitoring. Because PM has been associated with dysfunction in neural systems responsible for motor prediction and error monitoring,^9^ the CDT offers a promising behavioral measure for detecting altered motor awareness that may be specific to PM. Based on previous research on altered motor awareness in PwP,^15^ we hypothesized that (1) performance on the CDT would differ between healthy controls and PwP, and (2) among PwP, task performance would vary depending on the subtype of micrographia, with PwP with PM showing poorer performance due to impaired predictive and monitoring processes.

## Methods

### Participants

Twenty healthy individuals (73.6 ± 6.0 years old, 11 females) and 45 PwP undergoing rehabilitation (74.2 ± 6.8 years old, 24 females, Hoehn and Yahr scale stage 2.8 ± 0.9) were included in this study (Table 1). This study was performed in line with the principles of the Declaration of Helsinki. Approval was granted by the Ethics Committee at The University of Tokyo (approval number: KE23-35). Each participant provided written informed consent prior to participation.

**Table 1.** Demographic and clinical characteristics of the HC and PD groups.

|  | PD | HC |
| --- | --- | --- |
| Number | 45 | 20 |
| Age | 74.2 ± 6.8 | 73.6 ± 6.1 |
| Sex (F/M) | 24 / 21 | 11 / 9 |
| MMSE score, median (range) | 27.5 (21–30) | 29 (26–30) |
| Hoehn and Yahr stage, median (range) | 2.75 (1–4.5) | – |
| MDS-UPDRS Part III | 23.9 ± 13.5 | – |
| Disease duration | 6.7 ± 5.3 | – |
| Levodopa equivalent dose, mg/day | 760.4 ± 505.9 | – |

Mean ± SD values are shown, unless otherwise indicated. MMSE: Mini-Mental State
Examination; UPDRS: Unified Parkinson's Disease Rating Scale

The inclusion criteria for the healthy control (HC) group were as follows: (1) the ability to understand instructions; (2) absence of visual impairments or upper limb symptoms that could interfere with handwriting tasks; and (3) right-handedness. For the PwP group, the inclusion criteria were: (1) a definitive diagnosis of Parkinson’s disease based on the MDS-UPDRS ^17^; (2) age of 20 years or older; (3) if motor fluctuations (wearing off) were present, confirmation that participants were in the ON state at the time of the experiment; (4) absence of visual impairments or upper limb symptoms that could interfere with handwriting tasks; and (5) right-handedness.

Exclusion criteria for all participants included: (1) having a disease other than PD that affects the central nervous system; (2) presence of severe psychiatric symptoms such as hallucinations; (3) motor fluctuations not associated with medication timing; and (4) a sudden worsening of PD symptoms within one week prior to the experiment.

In this study, to match the age and sex distributions between the HC and PwP groups, we first recruited the PwP group and then set the inclusion criteria for the HC group based on those distributions.

### Experimental procedure

Participants sat at a table and first underwent a handwriting assessment, then performed the CDT. The entire experiment took approximately 30 minutes per participant.

### Handwriting

For the handwriting assessment, a DTK-1301 LCD tablet (13.3-inch, HD resolution 1920 × 1080 pixels; Wacom, Tokyo) was placed on a table and connected to a PC, and the coordinates of the handwriting produced with the accompanying digitizer pen were recorded. A sequence of ten model characters (“ma,” the Japanese hiragana character) was displayed in a horizontal row in the upper-left area of the tablet screen. Participants were asked to reproduce the same character ten times, writing them in a horizontal row to the right of the model characters while maintaining approximately 8 mm height and width for each character and preserving alignment (Figure 1). No time limit was imposed, and participants were instructed to write at a comfortable pace. Handwriting trajectories were recorded at a sampling frequency of 180 Hz with a coordinate resolution of approximately 0.01 mm.

**Figure 1.**
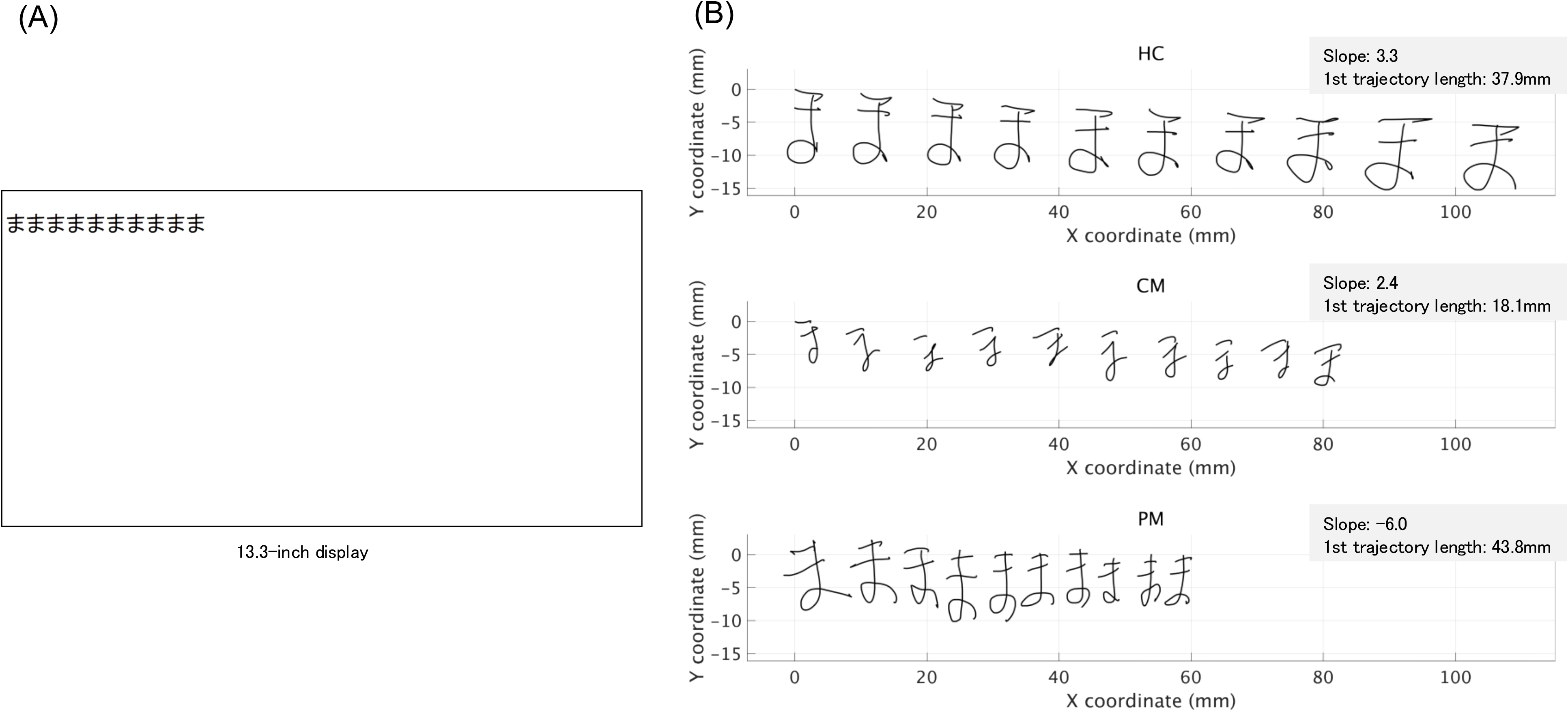
(A) Display used for the handwriting assessment task and representative handwriting samples from healthy individuals, CM patients, and PM patients. Ten example characters (“ma,” the Japanese hiragana character) were shown on the left side of the screen. Participants were instructed to reproduce the character ten times to the right, matching the displayed size. (B) Representative handwriting trajectories from a healthy control (HC), a patient with consistent micrographia (CM), and a patient with progressive micrographia (PM). The x- and y-coordinates indicate pen movement during successive reproductions of the character. Linear regression slopes (Slope) reflect changes in character size across repetitions, and the first trajectory length denotes the stroke length of the initial character.

In accordance with a previous study,^9^ participants were classified into three groups based on the trajectory length of each written character: a group without micrographia (w/o group), a group with CM, and a group with PM. The CM group was defined as cases in which the trajectory length of the first character was less than the mean minus 1.5 standard deviations of the HC group. To identify the PM group, each participant’s character trajectory lengths were first normalized by dividing the trajectory length of each character by that participant’s mean trajectory length across all ten characters. A linear regression was then performed on the normalized values across the writing sequence (characters 1–10), and the resulting slope was calculated. Participants whose slope was more than 1.5 standard deviations below the HC mean (i.e., a more negative value indicating a reduction in character size) were classified as having PM.

### Control detection task

The CDT ^16^ was used to assess awareness of motor control. This task has been employed to evaluate sensorimotor self-recognition in both healthy and clinical populations, including individuals with developmental coordination disorder.^18,19^ The CDT was conducted on a PC with a 15.6-inch LCD monitor (1920×1080 pixels, 60 Hz), keyboard, and mouse. Two white markers (one circle, one square; each approximately 6 × 6 mm) appeared on the screen. When participants moved the mouse, both markers moved, but the relationship between mouse movement and marker trajectory (i.e., direction) varied across trials.

Participants were asked to move a computer mouse for up to 10 seconds and judge which of two on-screen markers they believed they were controlling (Figure 2). One marker followed a prerecorded trajectory independent of the participant’s movement, whereas the other marker’s movement was generated by blending the participant’s real-time cursor motion with the prerecorded trajectory at ratios of 20%, 40%, 60%, 80%, or 100% (100% = pure participant-driven motion). If participants made a confident decision before 10 seconds elapsed, they could press the space bar to end the trial. When the trial ended—either after 10 seconds of upon keypress—both markers stopped. Numbers (“1” or “2”) then appeared next to the markers together with the prompt “Which marker do you feel you could control?” Participants responded verbally or via keypress, and the experimenter recorded the response.

**Figure 2.**
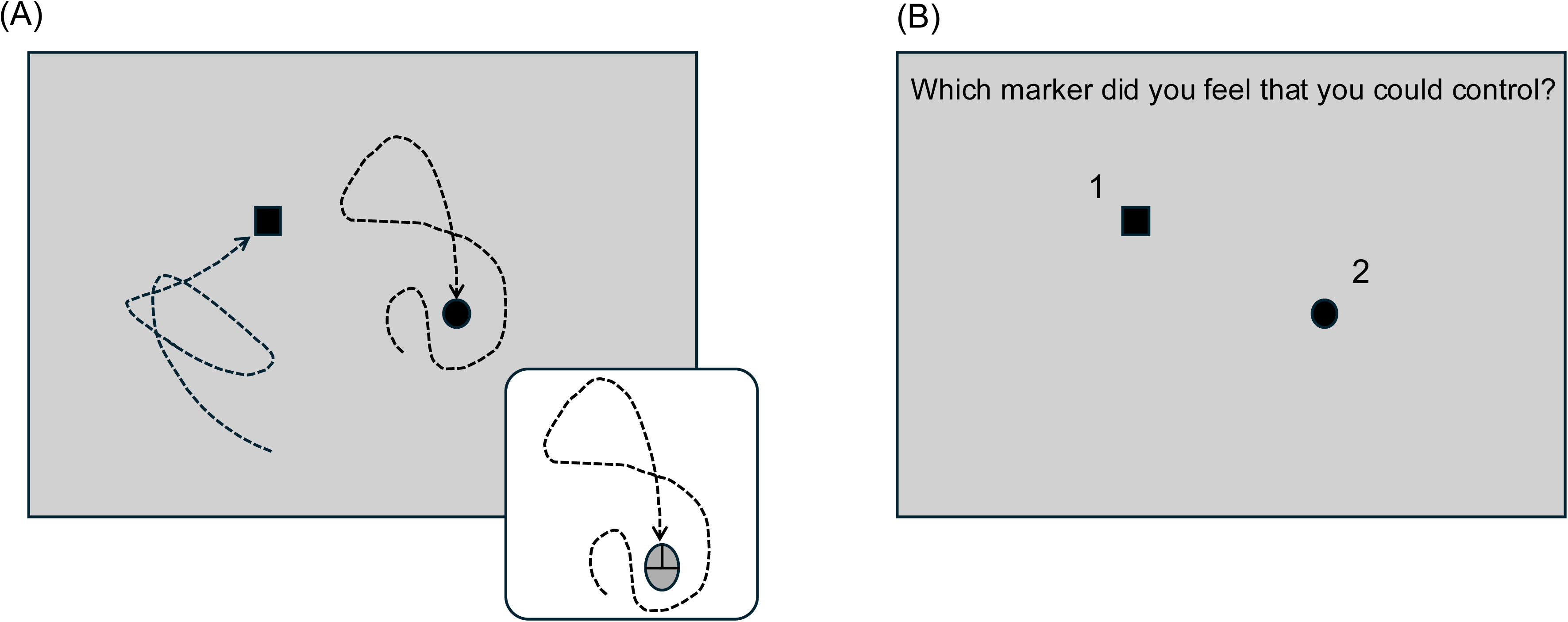
Representative trial displays from the control detection task. (A) Two markers (a circle and a square) moved on the screen in response to the participant’s mouse movement. (B) Participants judged which marker they were controlling based on the perceived correspondence between their movement and the marker trajectories.

A total of 30 trials were conducted in random order, with each blending condition appearing six times. Prior to the main task, participants completed six practice trials to familiarize themselves with the procedure. During the task, mouse trajectories were recorded continuously, and the total path length of each trajectory was calculated as an index of movement effort during the decision process. Reaction time was also recorded for each trial and defined as the interval between trial onset and the space-bar press by experimenter (or trial timeout).

### Clinical information

Patient clinical data was reviewed to obtain demographic and medical information. The collected data included age, sex, disease duration, severity of Parkinson’s disease, levodopa equivalent daily dose (LEDD), and results from neuropsychological assessments. Disease severity was evaluated based on the modified Hoehn and Yahr scale and MDS-UPDRS Part III. Neuropsychological assessments included the Mini Mental State Examination (MMSE) and the Frontal Assessment Battery (FAB).

## Statistical analysis

To compare patient characteristics across micrographia subtypes, linear mixed-effects models (LMMs) were used with CM, PM, and the interaction between CM and PM as fixed factors. LMMs were applied to continuous variables (age, disease duration, LEDD) and ordinal variables (MDS-UPDRS motor score, modified Hoehn and Yahr scale, MMSE, and FAB). One participant had missing MMSE and FAB data, which were handled automatically by the LMM using available cases.

Statistical analyses were performed using SPSS version 28 (IBM Corp., Armonk, NY, USA). For comparisons between the HC and PwP groups, the Mann–Whitney U test was applied, with Bonferroni-corrected post hoc tests where appropriate. Effect sizes (*r*) were calculated by dividing the standardized test statistic (*Z*) by the square root of the total number of observations. To confirm the adequacy of the sample size, a post hoc power calculation was performed using G*Power 3.1. Based on the observed effect size for CDT accuracy (Cohen’s *d* = 1.07), the achieved power was 0.96, indicating sufficient sensitivity for the HC–PwP comparison.

To examine relationships between handwriting performance and CDT outcomes across all participants (HCs and PwP), Spearman’s rank correlation coefficients were calculated. Specifically, correlations were assessed between the slope of handwriting character size across ten characters and CDT accuracy, CDT response time and mouse trajectory length during task responses.

For comparisons within the PwP group across micrographia subtypes (presence/absence of CM and PM), LMMs were used, with CDT accuracy, mouse trajectory length during task responses, and CDT response time entered as dependent variables. Fixed effects included CM and PM status, and random intercepts were specified for participants. Model assumptions were checked by examining residuals, which showed no substantial deviation from normality. The significance level was set at 5% for all analyses. Data are shown as mean ± SD unless otherwise indicated.

## Results

### Trajectory length of handwriting

In the HC group, the average handwriting trajectory length was 38.7 ± 6.8 mm for the first character, and 41.6 ± 7.9 mm for the tenth. In the PwP group, the corresponding values were 33.1 ± 8.9 mm and 30.4 ± 10.8 mm, respectively (Figure 3A). The average slope of character size across ten characters was 0.76 ± 1.2 in the HC group and-1.00 ± 3.4 in the PwP group.

**Figure 3.**
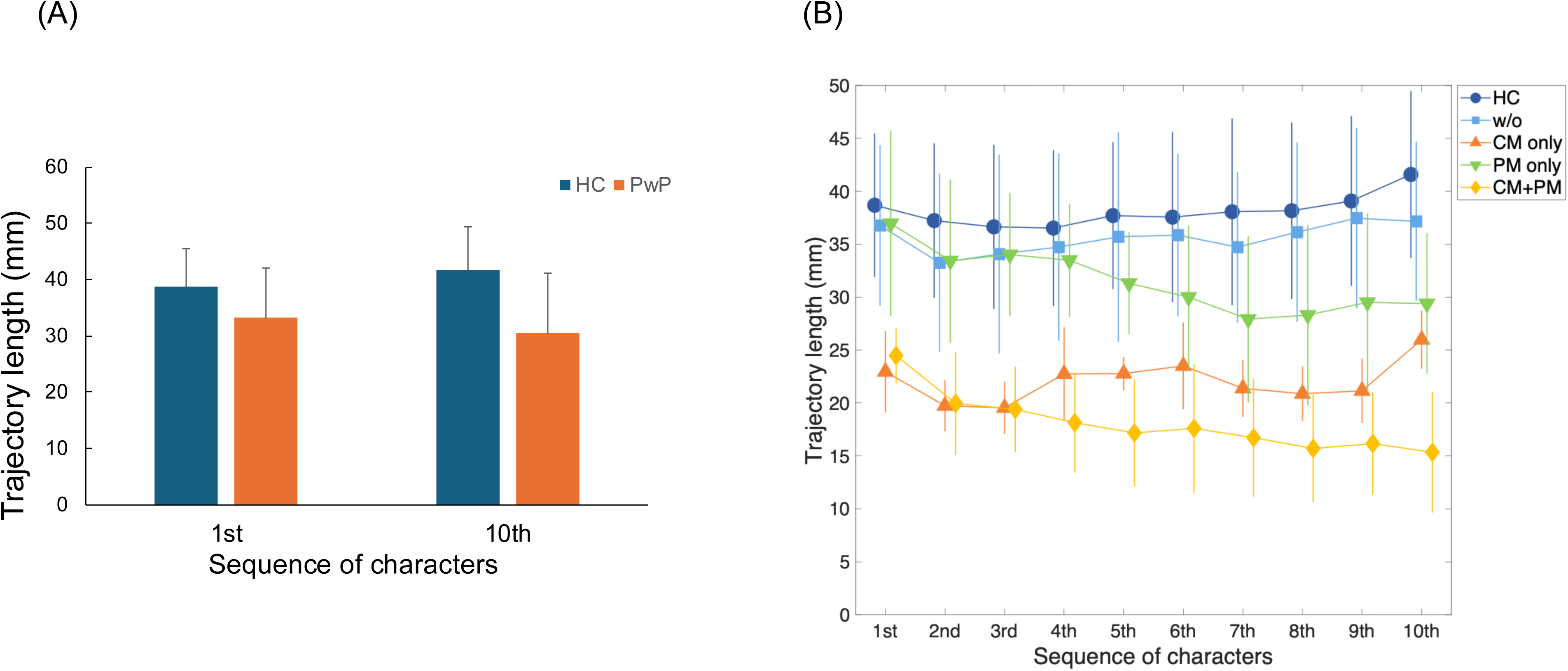
Trajectory length of handwriting. The mean values and standard deviations for the healthy control (HC) group and the Parkinson’s disease (PwP) group are shown. (A) Comparison of trajectory length between the 1st and 10th character sequences. (B) Changes in trajectory length across all 10 character sequences in each group. Error bars indicate standard deviations.

Based on HC data, PwP handwriting was classified according to four groups: w/o (n = 24), CM only (n = 4), PM only (n = 8), and CM + PM (n = 9). Demographic and clinical characteristics are shown in Supplemental Table S1. These characteristics were compared across micrographia subtypes using linear mixed-effects models with CM, PM, and the interaction between CM and PM as fixed factors. The models revealed significant main effects of PM on MMSE (*F*(1, 40) = 5.81, *p* = 0.02) and FAB (*F*(1, 40) = 4.78, *p* = 0.04).

No significant main effects of CM or interaction between CM and PM were observed for either variable (*p* > 0.05). In contrast, age, disease duration, HY, UPDRS, and LEDD showed no significant main or interaction effects (all *p* > 0.05).

### Accuracy, response time and trajectory length in the CDT

Mean CDT accuracy across blending conditions was 88.8 ± 7.1% in HCs and 74.4 ± 17.7% in PwP, a significant group difference (effect size *r* = 0.43, *p* < 0.01; Figure 4A). The percentage of correct responses by control level in HCs was 70.0% (control level 20%), 87.5% (40%), 95.0% (60%), 93.3% (80%), and 98.3% (100%), and in PwP was 57.8%, 68.1%, 77.8%, 84.1%, and 84.4%, respectively. Significant correlations were observed between the handwriting slope of character size across ten characters and CDT accuracy (*rs* = 0.34, 95% CI [0.09, 0.54], *p* < 0.01), as well as between the handwriting slope and CDT response time (*rs* =-0.32, 95% CI [-0.52,-0.07], *p* = 0.01). In contrast, no significant correlation was found between handwriting slope and mouse trajectory length during task responses (*rs* = 0.02, 95% CI [-0.23, 0.41], *p* = 0.27, *p* = 0.87).

**Figure 4.**
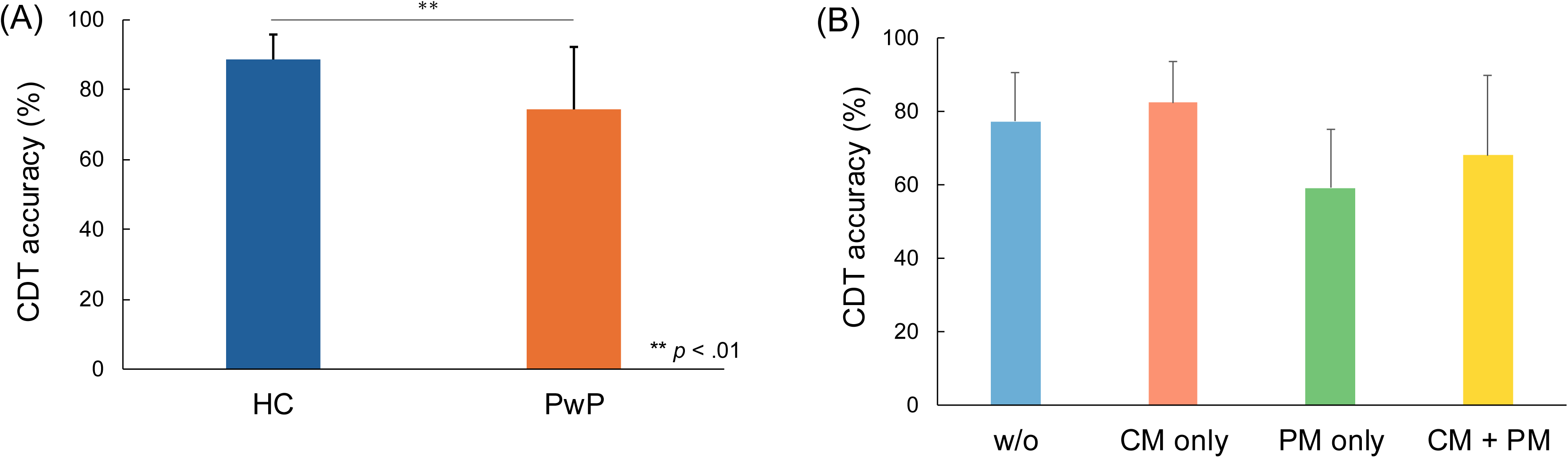
Accuracy in the control detection task (CDT). (A) Average CDT accuracy was significantly higher in the healthy control (HC) group compared to the Parkinson’s disease (PwP) group (*p* <.01). (B) Comparison of accuracy based on micrographia classification. The linear mixed-effects model revealed a significant main effect of progressive micrographia (*p* <.01). Error bars indicate standard deviations.

Among the micrographia subtypes, mean CDT accuracy across blending conditions was 77.3 ± 13.4% in the w/o group, 82.5 ± 11.0% in the CM only group, 59.2 ± 16.0% in the PM only group, and 68.1 ± 21.6% in the CM + PM group (Figure 4B and 4C). The linear mixed-effects model revealed a significant main effect of PM (*F*(1,41) = 9.59, *p* < 0.01). No significant main effect of CM or interaction between CM and PM was observed.

The average response time per trial, was 6.74 ± 1.60 s in the w/o group, 6.08 ± 0.91 s in the CM only group, 8.48 ± 1.89 s in the PM only group, and 8.37 ± 2.17 s in the CM + PM group. The linear mixed-effects model revealed a significant main effect of PM (*F*(1,41) = 8.49, *p* < 0.01). No significant main effect of CM or interaction between CM and PM was observed.

The average mouse movement trajectory length per trial, reflecting movement effort, was 927 ± 805 mm in the w/o group, 507 ± 348 mm in the CM only group, 1612 ± 1547 mm in the PM only group, and 561 ± 284 mm in the CM + PM group. The linear mixed-effects model revealed no significant main effects of CM or PM, and no significant interaction between CM and PM (*p* > 0.05).

## Discussion

This study examined whether altered awareness of motor control is linked to micrographia in PwP, using the CDT to assess self–other attribution of motor actions in PwP and healthy controls. Our findings revealed significantly lower CDT accuracy in PwP, particularly among those classified as having PM. In addition, correlation analyses demonstrated a relationship between the severity of micrographia, indexed by the handwriting slope, and CDT accuracy. To our knowledge, this is the first study to systematically evaluate sensorimotor awareness across micrographia subtypes in Parkinson’s disease using the CDT, providing a novel behavioral perspective on the mechanisms underlying handwriting impairments. While previous research has primarily addressed general handwriting deficits, our results highlight that PM is associated with marked disruptions in predictive and monitoring mechanisms of motor control.

The CDT requires participants to operate a mouse under visual feedback and identify which of two stimuli they are controlling, necessitating a comparison of predicted motor outcomes with actual sensory feedback. Thus, reduced performance in PwP likely reflects disruptions in neural mechanisms underlying action prediction and monitoring. This interpretation aligns with a previous study showing that children with developmental coordination disorder (DCD) exhibit reduced CDT performance.^19^ Importantly, the results of our present report support the same interpretation in an adult population: healthy older adults of comparable age to the PwP exhibited better performance on the CDT. This suggests that reduced CDT accuracy cannot be attributed to age-related decline, but instead reflects disease-specific deficits in prediction–feedback integration. Although the motor impairments in PwP and DCD differ, both groups may share difficulties in using feedback to adjust ongoing actions, possibly reflecting an imbalance between feedforward and feedback control strategies. Generally, feedforward mechanisms predominate in rapid, goal-directed movements,^20^ while feedback processes are crucial during early learning or extended-duration tasks.^21,22^ Because handwriting also relies on continuous integration of prediction and feedback, these findings raise the possibility that, especially in PM, altered feedback-based correction contributes to handwriting difficulties, potentially reflecting broader challenges in how feedback is used during motor learning. Furthermore, handwriting slope was significantly correlated with CDT accuracy across participants, suggesting that micrographia-related impairments in sensorimotor awareness may be captured not only by categorical subtyping but also along a continuous dimension. Notably, response time was also significantly prolonged in individuals with PM, indicating that reduced CDT accuracy in this group may reflect increased task difficulty and feedback-based decisional load rather than a simple error in control attribution.

Analysis of clinical features across micrographia subtypes showed that the PM group had greater motor severity, as indicated by a higher modified Hoehn and Yahr scale score than in the w/o group. This difference suggests an association between overall disease progression and the manifestation of PM. However, the trajectory length of mouse movement (an index of movement effort, reflected in the distance traveled during the decision process) during the CDT did not differ significantly among subtypes, indicating that reduced CDT accuracy in the PM group is unlikely to be explained solely by motor execution deficits. Taken together, these findings suggest that the deficit in PM more plausibly lies in the perception or processing of movement, supporting the interpretation that altered motor awareness—beyond pure motor output—underlies the observed performance deficit. Moreover, the PM group’s lower MMSE scores raise the possibility that cognitive decline or cortical dysfunction may contribute to PM.^23^ To our knowledge, no studies have directly investigated CDT performance in Alzheimer’s disease or other dementias. Nevertheless, recent work has demonstrated reduced handwriting fluency in patients with Alzheimer’s disease,^24^ albeit with patterns distinct from those in Parkinson’s disease, suggesting that cognitive decline can also impact handwriting-related functions. Further research comparing PwP with other clinical populations is warranted to clarify these relationships.

Consistent with the notion that deficits in PM are more likely to arise from altered perception or processing rather than from motor execution itself, previous neurophysiological studies have demonstrated abnormalities in brain networks involved in online visual processing and error monitoring in patients with PM.^9^ Other research has shown that visual feedback during handwriting plays a role in monitoring spatial characteristics and character size.^25^ Furthermore, studies in children have shown that when visual feedback of their handwriting is blocked, character size increases, likely due to a compensatory shift toward greater reliance on somatosensory feedback.^26^ Collectively, these findings suggest that individuals with PM may exhibit altered processing of visual feedback or may be unable to enhance somatosensory gain as a compensatory strategy.

Although a significant relationship between handwriting slope and CDT accuracy was observed across participants, inspection of individual data revealed heterogeneity within the CM + PM group. Specifically, a small subset of individuals showed relatively preserved CDT accuracy despite pronounced handwriting slopes (Figure 5B; cases located in the lower-right area). When these three heterogeneous cases were excluded in an exploratory analysis, the correlation coefficient increased to *rs* = 0.47, suggesting that the overall association may be attenuated by a limited number of atypical data points. Although limited in number, this dissociation suggests the presence of heterogeneous patterns in the relationship between handwriting impairment and sensorimotor awareness, even within the same micrographia subtype. One possible explanation for the observed dissociation may be the influence of Parkinson’s disease phenotypes, which may differ in terms of motor subtype and cognitive involvement. ^27,28^ In addition, individual differences in cognitive strategy, attentional allocation, or reliance on visual versus proprioceptive feedback during the CDT, as well as variability in compensatory neural mechanisms or medication responsiveness,^29,30^ may further contribute to this heterogeneity. Since these interpretations are speculative, further investigations using neuroimaging and sensorimotor modeling approaches are warranted.

**Figure 5.**
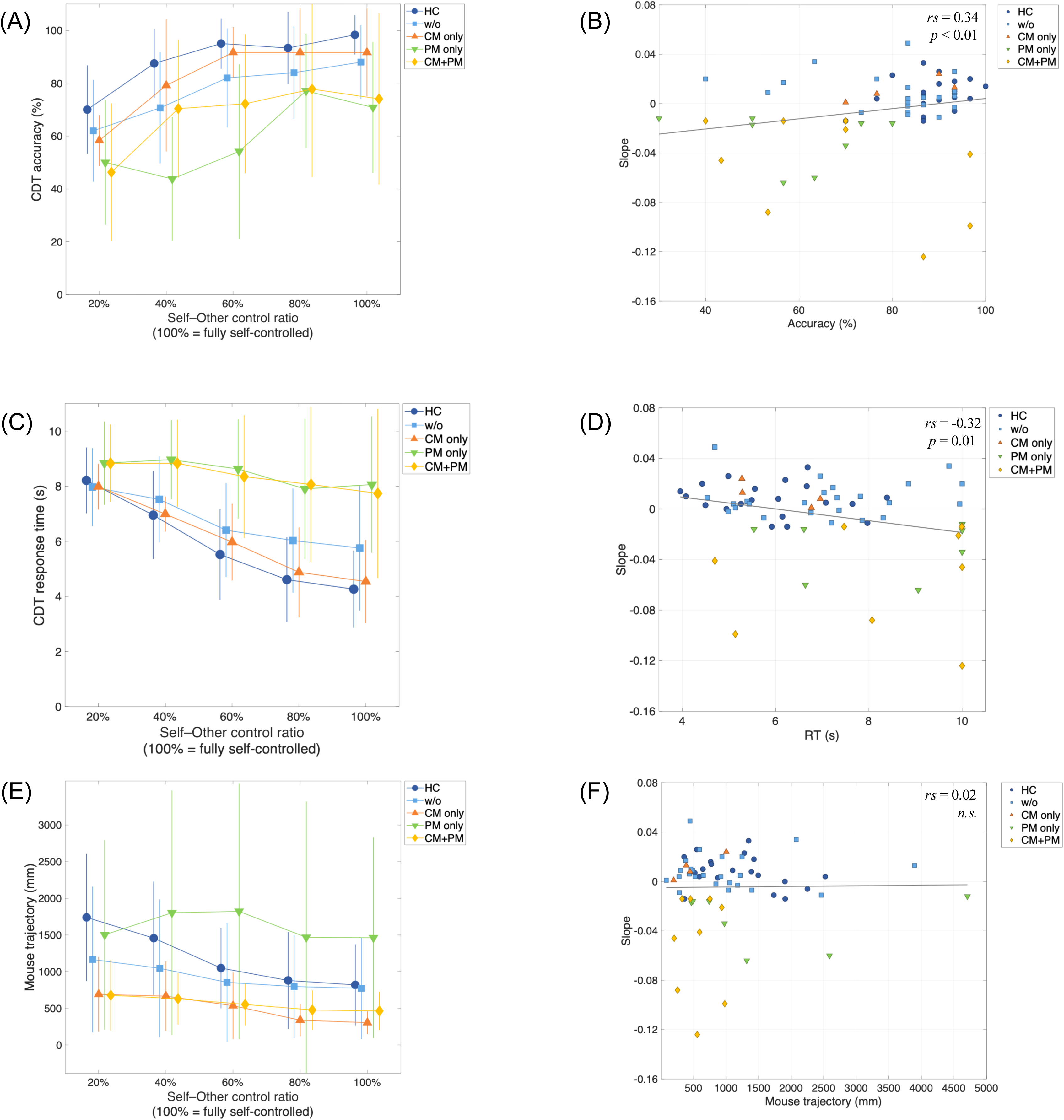
Control detection task performance and correlation analysis (A) Mean CDT accuracy (%) across self–other control ratios (20–100%, where 100% indicates fully self-controlled movement) for healthy controls (HC) and Parkinson’s disease subgroups (w/o, CM only, PM only, and CM + PM). Error bars indicate standard deviation. (B) Relationship between handwriting slope of character size across ten characters and CDT accuracy. Each point represents an individual participant. A significant positive correlation was observed (*p* <.01). (C) Mean CDT response time across self–other control ratios for each group. (D) Relationship between handwriting slope and CDT response time, showing a significant negative correlation (*p* = 0.01). (E) Mean mouse movement trajectory length per trial across self–other control ratios for each group. (F) Relationship between handwriting slope and mouse movement trajectory length, showing no significant correlation. Data points in panels (B), (D), and (F) are color-coded according to group (HC, w/o, CM only, PM only, and CM + PM). Solid lines in the scatter plots represent linear regression fits for visualization purposes only. In the left panels, error bars represent standard deviations.

From a clinical perspective, the present findings offer new insights into rehabilitation strategies for handwriting impairments in PwP. If sensorimotor mismatch processes are involved in the pathophysiology of PM, interventions that focus solely on motor output may be insufficient. Training approaches that encourage patients to consciously compare predicted outcomes with actual sensory feedback, combined with external feedback on performance accuracy, may therefore be more effective. For instance, approaches that encourage PwP to consciously compare predicted outcomes and actual sensory feedback, combined with external feedback on performance accuracy could be adapted as training tasks. In PwP, adaptive adjustments do not typically occur when visual feedback of handwriting size is experimentally altered (e.g., making the written trace appear smaller does not induce compensatory enlargement of the actual writing).^31^ However, successful learning has been observed in visuomotor transformation tasks such as prism adaptation.^32^ These observations suggest that rehabilitation may be more effective when tasks are designed to facilitate attention to feedback—for example, by slowing down movement speed while emphasizing performance monitoring. Future research should focus on developing and validating such rehabilitation strategies.

This study has several limitations. First, because the CDT reflects a combination of processes such as motor prediction, sensory feedback integration, and error monitoring, complementary tasks were not included to disentangle these components or provide converging evidence. Second, although the CDT engages distributed neural circuits, we did not combine behavioral performance with neuroimaging, which would have allowed identification of the key networks underlying control detection in PwP and informed potential intervention strategies. Third, while we classified micrographia subtypes, some overlap among groups existed, which may reflect disease progression or additional unmeasured factors. Longitudinal, multimodal, and more detailed investigations will be needed to address these issues.

## Conclusion

Our findings demonstrate that individuals with Parkinson’s disease have impaired motor awareness compared to healthy controls, with those exhibiting PM showing the most pronounced deficits in sensorimotor self-recognition. Disrupted prediction–feedback integration likely underlies PM and reflects broader neural dysfunction related to handwriting impairment in PwP. These results highlight the promise of motor-awareness–based rehabilitation as a novel approach to addressing micrographia.

## Acknowledgments

The authors would like to thank Ryoma Aoki, Takuya Iwamoto, Keisuke Shiraishi, Natsumi Takahashi, Kazuki Yoshida, and Akika Yoshimoto, rehabilitation therapists at the Department of Rehabilitation, Noborito Neurology Clinic, for their assistance with data collection.

## Author Contributions

Hiroyuki Hamada: Conceptualization, Data curation, Formal analysis, Funding acquisition, Investigation, Methodology, Project administration, Resources, Software, Supervision, Visualization, Writing – original draft, Writing – review & editing

Kyohei Mikami: Conceptualization, Data curation, Formal analysis, Investigation, Writing – review & editing

Wen Wen: Conceptualization, Software, Formal analysis, Writing – review & editing Yoshihiro Itaguchi: Conceptualization, Software, Formal analysis, Visualization, Writing – review & editing

Atsushi Yamashita: Conceptualization, Formal analysis, Writing – review & editing

Qi An: Conceptualization, Formal analysis, Funding acquisition, Writing – review & editing

## Statements and declarations Ethical considerations

This study was performed in line with the principles of the Declaration of Helsinki. Approval was granted by the Ethics Committee at The University of Tokyo (approval number: KE23-35).

## Consent to participate

Each participant provided written informed consent prior to participation.

## Consent for publication

Not applicable.

## Data availability statement

The data presented in this study are available from the corresponding author upon reasonable request, subject to institutional and ethical approval.

